# Zebrafish airinemes optimize their shape between ballistic and diffusive search

**DOI:** 10.1101/2021.10.24.465630

**Authors:** Sohyeon Park, Hyunjoong Kim, Yi Wang, Dae Seok Eom, Jun Allard

## Abstract

In addition to diffusive signals, cells in tissue also communicate via long, thin cellular protrusions, such as airinemes in zebrafish. Before establishing communication, cellular protrusions must find their target cell. Here we demonstrate that the shape of airinemes in zebrafish are consistent with a finite persistent random walk model. The probability of contacting the target cell is maximized for a balance between ballistic search (straight) and diffusive search (highly curved, random). We find that the curvature of airinemes in zebrafish, extracted from live cell microscopy, is approximately the same value as the optimum in the simple persistent random walk model. We also explore the ability of the target cell to infer direction of the airineme’s source, finding that there is a theoretical trade-off between search optimality and directional information. This provides a framework to characterize the shape, and performance objectives, of non-canonical cellular protrusions in general.

## 1 Introduction

The question of optimal search — given a spatio-temporal process, what parameters allow a searcher to find its target with greatest success? — arises in many biological contexts for a variety of spatio-temporal processes. Examples of relevant processes include searchers moving by diffusion or random walks [1, 2], Levy walks [3], and ballistic motion (straight trajectories [5], which, e.g., arises in chromosome search by microtubules [6, 7]), and combinations of these [8]. Another type of motion is the persistent random walk, which has intermediate properties between diffusion and ballistic motion, and has been studied in continuous space [9–11] and on a lattice [4]. For all the above processes, optimality depends on parameters of the searcher (e.g., whether searchers operate individually or many in parallel [12, 13]), the target(s), and the environment [5, 14].

One example of a biological search process arises during organismal development, when cells must establish long-range communication. Some of this communication occurs by diffusing molecules [15–18] like morphogens. However, recently, an alternative cell-cell communication mechanism has been revealed to be long, thin cellular protrusions extending 10s-100s of micrometers [19–24]. These include cytonemes [25], nanotubes [26], and airinemes in zebrafish [27–30], shown in Fig. 1A. One of the difficulties delaying their discovery and characterization is their thin, sub-optical width, and the fact that they only form at specific stages of development [19–21].

**Figure 1:**
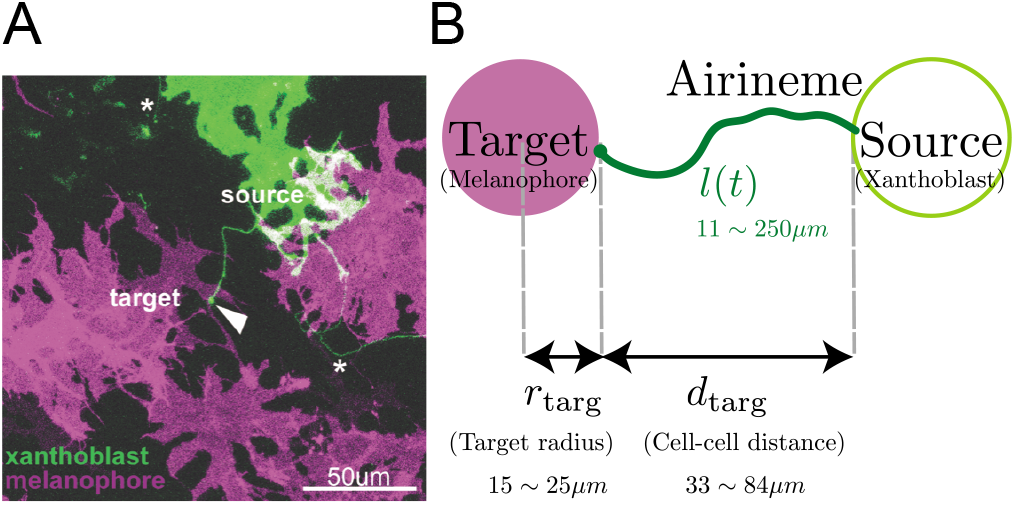
**(A)** Multiple airinemes extend from xanthoblast (undifferentiated yellow pigment cell, green). Airineme makes successful contact (arrowhead) with melanophore cell (pigment cell, purple). Asterisks indicate airinemes from other sources. Scale bar: 50*μm*. **(B)** Model schematic. A single airineme extends from the source (right, green circle) and searches for the target cell (left, purple circle). Target cell has radius *r*_targ_, and has distance *d*_targ_ away from the origin. The airineme’s contour length at time *t* is *l*(*t*).

For diffusing cell-cell signals, dynamics are characterized by a diffusion coefficient. In contrast, cellular protrusions require more parameters to describe, e.g., a velocity and angular diffusion, or equivalently a curvature persistence length. Here, we ask, what are these parameters, and what determines their values? We focus on airinemes where quantitative details have been measured [29, 30]. We find that airineme shape is most consistent with a finite-length persistent random walk model, and that the parameters of this model exhibit an optimum for minimizing the probability of finding a target (or, equivalently, the mean number of attempts). We compare this with another performance objective, the ability for the airineme to provide a directional cue to the target cell, and find that there is a theoretical trade-off between these two objectives. This work provides an example where a readily-observable optimum appears to be obtained by a biological system.

## 2 Results

### 2.1 Airinemes are consistent with a finite persistent random walk model

We examined time-lapse live cell image data as described in [29, 30]. We confirmed that the time series and the final-state are similar (Fig. S1), meaning the shape of an airineme does not change much throughout its extension. This allows us to consider only the fully extended airineme and infer the dynamics, assuming airnemes extend with constant velocity *v* = 4.5*μm*/min [30]. This removes artifacts like, e.g., microscope stage drift, and drastically simplifies the analysis. We manually identified and discretized 70 airinemes into 5596 position vectors *r*(*t*), and from these, computed the mean squared displacement (MSD) 〈*r*^2^〉. Random walks satisfy 〈*r*^2^〉 = 4*Dt*. However, the observed MSD, shown in Fig. 2A, has a best-fit exponent of 1.55 (90% CI in [1.50,1.61]), and indeed it appears that a single exponent is not appropriate across orders of magnitude. We therefore reject the simple random walk description.

**Figure 2:**
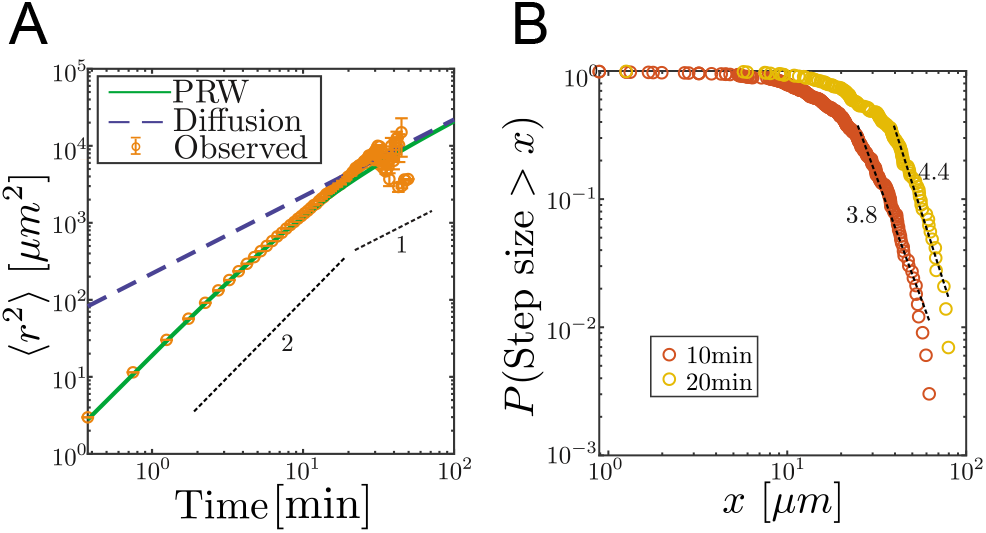
Airinemes are not consistent with simple random walk or Levy-type models. **(A)** Mean squared displacement (MSD) of airinemes. Airineme experimental MSDs have exponent ≈ 2 (corresponding to slope in log-log plot) up to ≈ 30 minutes, arguing against a simple random walk model. *N* = 56 airinemes **(B)** Step length complementary cumulative distribution function (CCDF). Levy models in 2d are characterized by CCDF tails with an exponent between 1 and 3. However, for two different time sampling intervals (10 and 20 minutes, from *N* = 56 airinemes), the tails of the distribution fit to an exponent greater than 3, and a continued downward curvature instead of a power-law, arguing against Levy models.

Next, we consider Levy-type models such as those that have been used to describe animal optimal foraging [31] and T cell migration [3]. These processes have a step size distribution whose tail has exponent between 1 and 3 in 2d [3, 31]. We revisit the time-series data (i.e., here we do not use the final state approximation) and compute a step length complementary cumulative distribution function (CCDF). For two time interval choices, shown in Fig. 2B, the best fit CCDF exponents are greater than 3 (for 10 min, exponent is 3.81 with 90% CI in [3.68,3.95]; for 20 min, exponent is 4.40 with 90% CI in [4.18,4.61]). Indeed, the CCDFs at two different time sampling intervals have continued downward curvature, indicating that a power-law description is inappropriate. We thus conclude that the process is not consistent with Levy-type models.

Finally, we consider a finite-length persistent random walk (PRW). In this model, the tip of the airineme moves at constant speed *v*, while the direction undergoes random changes with parameter *D_θ_*, the angular diffusion coefficient. This parameter has units inverse minutes, and roughly corresponds to the “curviness” of the path. The dynamics are governed by Eqs. 2-4.

The airineme MSD fits the prediction of the PRW model, in Eq. 5, up to time point around 10 minutes. Above this time, the PRW model is consistent with the data, although the low number of long airinemes in our data precludes a stronger conclusion. (Below in Fig. 4, we also check the autocorrelation function and further confirm consistency with the PRW model.) Taken together, the data favor the PRW model, which we use in the following analysis.

We also assume airinemes operate independently, as there is no evidence of airinemes communicating with each other during the search process. Furthermore, airinemes are generated at approximately 0.15 airinemes per cell per hour. Thus the mean time between airineme initiations is ≈ 400 min, much larger than the time each airineme extends, which is 56min. Note that many airinemes emanating from the same source cell may exist simultaneously, but most of the time only one airinemes is extending. Also, while the tissue surface is crowded, the airineme tips (which are transported by macrophages [30]) appear unrestricted in their motion on the 2d surface, passing over or under other cells unimpeded [30]. We therefore do not consider obstacles in our model (although these have been studied in other PRW contexts [9, 11, 34]).

The target cell is modeled as a circle of radius *r*_targ_ = 15 – 25*μm* [30], separated from the source of the airineme by a distance *d*_targ_ ≈ 50*μm*, as shown in Fig. 1B. Including the position and size of the target, the model has 5 parameters, all of which have been measured (see Table S1 and [30, 46]) except for *D_θ_*.

### 2.2 Contact probability is maximized for a balance between ballistic and diffusive search

We performed simulations of the PRW model, testing different angular diffusion values for different values of given cell-to-cell distance and target cell radius, shown in Fig. 3AB. For each parameter set, we measured the proportion of simulations that contacted the target. Equivalently, we plot its inverse, the mean number of attempts, in Fig. S2.

**Figure 3:**
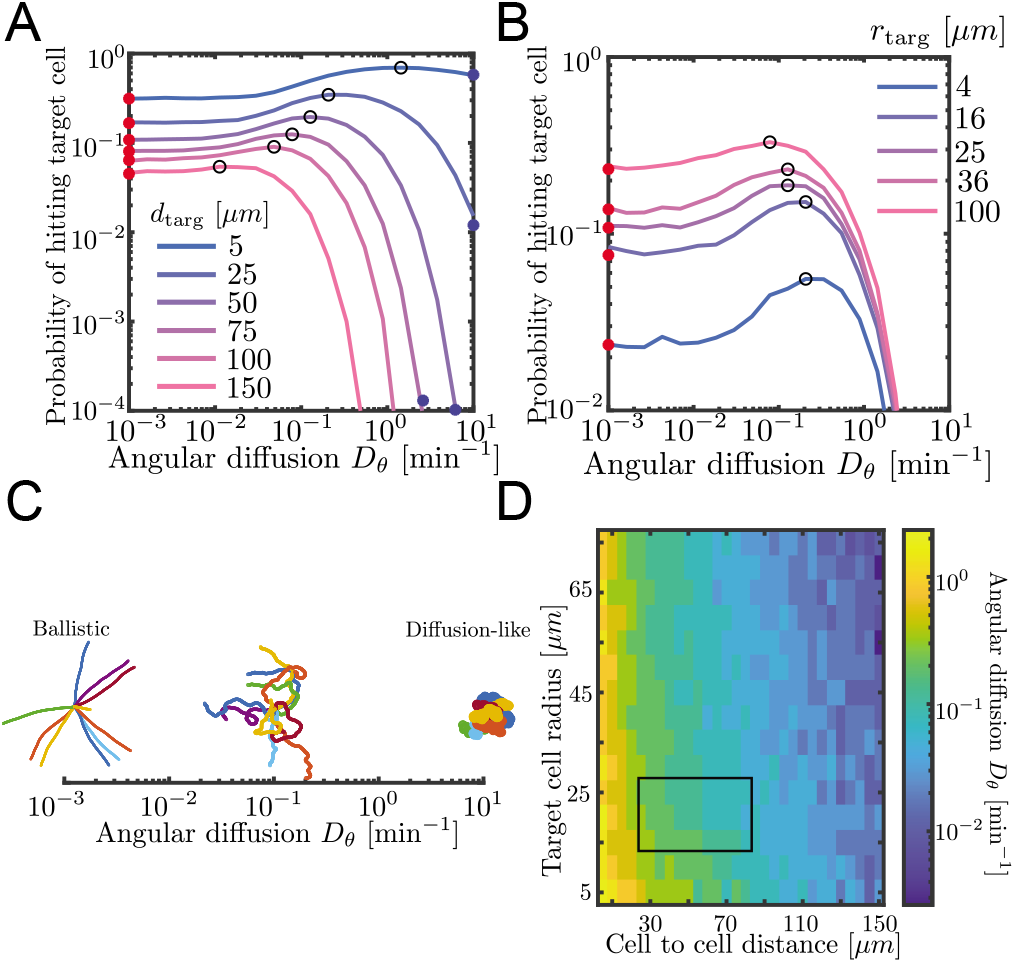
Probability to contact a target cell is maximized by a balance between ballistic search and diffusion-like search. **(A,B)** Simulated airineme target search for various *D_θ_*. The value of *D_θ_* allow the airineme to hit the target cell with largest probability of contact are shown in black open circles. Theoretical values for ballistic limit from (6) are shown as red circles. **(A)** Varying distances between the target and the source cell, while fixing the target cell radius at 25*μm*. We validate simulation results with survival probability PDE at high *D_θ_* from Eq. 7, shown in blue circles. **(B)** Varying target cell radii while fixing the distance between the target and the source. For the biologically relevant parameter (*d*_targ_ ~ 50*μm*), optimal angular diffusion is around *D_θ_* ~ 0.18 min^−1^. **(C)** Qualitative behavior of airineme target search depends on *D_θ_*. **(D)** Optimal *D_θ_* for a larger range of cell-to-cell distances and target cell radii. Rectangular region shows biologically relevant parameters.

**Figure 4:**
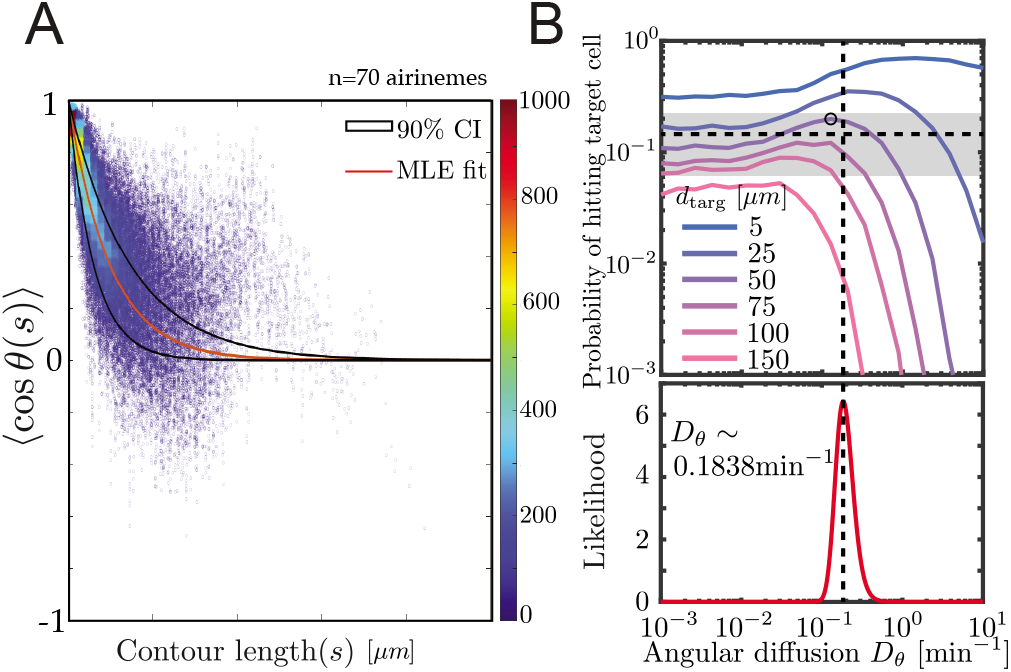
Experimental airineme curvature agrees with the optimal curvature. **(A)** Orientation autocorrelation function. We measure tangent angles cos (*θ*) at 5596 points along 70 airinemes, and then compute the likelihood function (B, bottom) of *D_θ_* fit to Eq. (1). Best-fit curve is shown as red with a 90 percent confidence interval shown in black. Bottom, we find the best fit airineme curvature from maximum likelihood estimation is *D_θ_* = 0.1838min^−1^. We find this value is similar to the *D_θ_* that optimizes contact probability for the biologically relevant target cell distance *d*_targ_ = 50*μm* (top). The experimentally-observed probability of contact per airineme, center estimate (horizontal dashed line) and 90% confidence interval (gray area), also shown.

We find that there exists an optimal angular diffusion coefficient that maximizes the chance to contact the target cell. The optimal value balances between ballistic and diffusion-like search. This has been previously shown for infinite, on-lattice persistent random walks [4], and we heuristically understand it as follows. When *D_θ_* is small (Fig. 3C left), airinemes are straight and therefore move outward a large distance, which is favorable for finding distant targets. However, straight airinemes easily miss targets. On the other hand, for *D_θ_* large (Fig. 3C right), the airineme executes a random walk. Random walks are locally thorough, so do not miss nearby targets, but the search rarely travels far. Thus, if the target cell is small or close, a diffusion-like search process is favored, but if the target cell is far or large, then a ballistic search is favored. We confirm this in Fig. 3D, where we plot optimal *D_θ_* over a large range of target cell radii and cell to cell distances. For the biologically relevant parameters (rectangular region in Fig. 3D), a balance between ballistic and diffusion-like is optimal.

### 2.3 Experimental airineme curvature is approximately optimal

In order to estimate the missing parameter *D_θ_*, we use the angular autocorrelation function

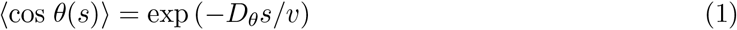

shown in Fig. 4. We performed manual image analysis and maximum likelihood estimation to fit Eq. 1, along with model convolution (Fig. S3, [35], Methods) to estimate uncertainty.

We find the maximum likelihood estimated persistence length of 12.24*μm*, corresponding to an angular diffusion *D_θ_* = 0.184min^−1^. Surprisingly, as shown in Fig. 4B, this value matches with our simulated optimal angular diffusion value for the biologically relevant parameter values. Moreover, in the experimental data, we find that the proportion making successful contact with target cells is *P*_Contact_ = 0.15 (horizontal dashed line), from *N* = 49 airinemes with a 90% confidence interval in [0.06, 0.24] (gray box in Fig. 4B). This also agrees surprisingly well with the model prediction *p*_contact_ ≈ 0.185.

### 2.4 Directional information at the target cell

In some models of zebrafish pattern formation, the target cells receive directional information from source cells [19, 28], i.e., the target cell must determine where the source cell is, relative to the target’s current position. We explore the hypothesis that airineme contact itself could provide directional information, since the location on the target cell at which the airineme contacts, *θ*_contact_ as shown in Fig. 5A, is correlated with the direction to the source cell. Analogous directional sensing is possible by diffusive signals, where physical limits have been computed in a variety of situations [1, 36].

**Figure 5:**
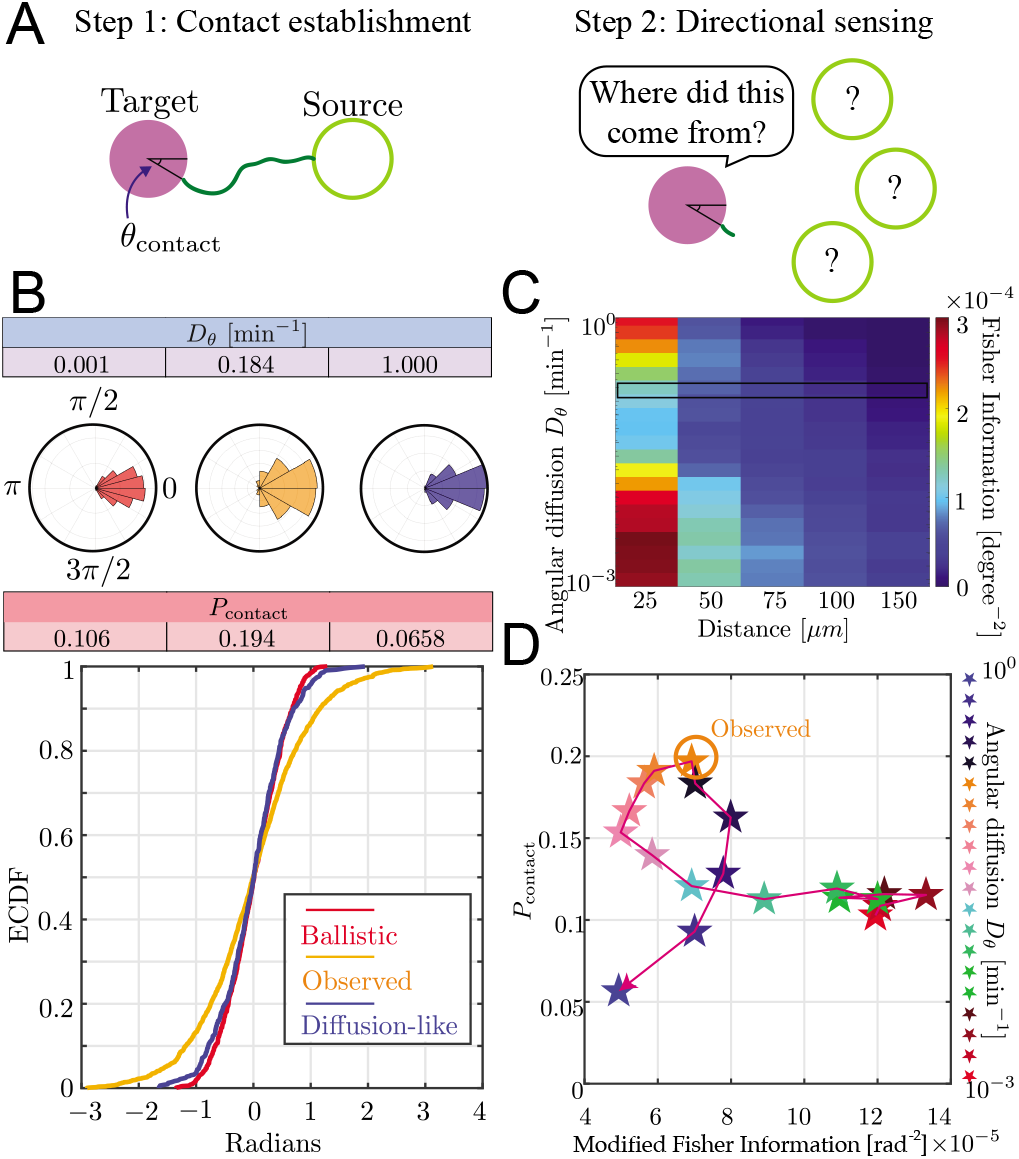
Trade-off between airineme directional sensing information and the probability of contacting the target cell. **(A)** Given that the source cell is located at *θ* = 0, the distribution of angles at which the airineme contacts the target cell. **(B)** Given a source cell is located at *θ* = 0, the distribution of angles at which the airineme contacts the target cell. Angle distributions with higher variance indicate poorer directional sensing. Three angular diffusion values (near ballistic limit, experimentally observed, and near diffusion-limit) are shown. **(C)** Directional sensing accuracy, quantified using the modified Fisher Information (FI) Eq. (9), for different source-to-target distances and different ranges of angular diffusion values are tested, while fixing target cell radius at 25*μm*. Black rectangular region shows the FI values for the observed airineme curvature. **(D)** Relationship between the contact probability *P*_contact_ and FI for a range of *D_θ_* (increasing with direction of arrow).

**Table 1:**
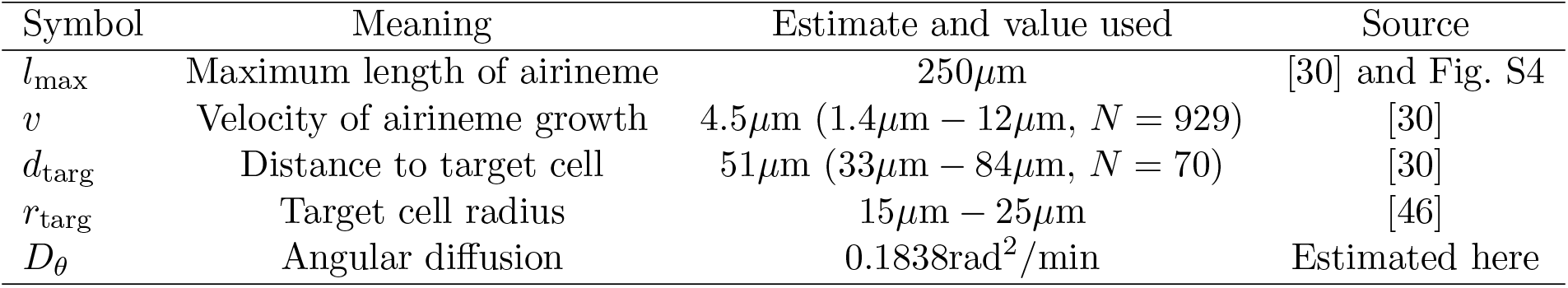
Parameter values.

We examined the contact angle distribution on the target cell. In Fig. 5B, we show this distribution for three values of *D_θ_*: low (ballistic), high (diffusion-like), and the observed value we found above. The source cell is placed at *θ*_origin_ = 0 without loss of generality. We show the distribution of contact angles on the target cell *p*(*θ*_contact_|*θ*_origin_ = 0) as both a radial histogram (top) and cumulative distribution (bottom). Interestingly, we find that the observed airineme parameters lead to a wide distribution of contact angles, compared to both ballistic or diffusion-like airinemes.

To quantify the ability of the target cell to sense the direction of the source by arrival angle of a single airineme, we use the Fisher Information [37] metric, modified to take into account the probabilistic number of airinemes that a target cell receives, using Eq. 9. Using this measure, we find that there is a rough trade-off between the airineme’s target contact success and the target cell’s directional sensing, as shown in Fig. 5D. Heuristically, this is because the two objectives prefer opposite variances. To maximize contact probability, variance should be maximal, taking full advantage of the surface of the target. On the other hand, to maximize directional information, the variance of contact angle should be minimized.

Inspecting both contact probability and directional information in Fig. 5D, we find that the experimentally observed *D_θ_* is at a point where either increasing or decreasing its value would suffer one or the other objectives, a property known as Pareto optimality [38, 39]. Note that this is also the *D_θ_* value that maximizes search success, so the data is consistent with either conclusion that the curvature is optimized for search, or it is optimized to balance search and directional information. In other words, in the case of zebrafish airinemes, there is no evidence that the shape of these protrusions sacrifices the goal of optimal search in order to achieve increased directional signaling.

To compare with experimental observations, we attempted to measure the contact angle distribution of airinemes in contact with target cells. This is complicated by the highly non-circular shape of these cells, so we approximate the angle by connecting 3 points: the point on the source cell from which the airineme begins, the center of the nucleus of the target, and the point on the surface of the target where the airineme makes contact, as shown in Fig. S5. We find a modified Fisher Information of 5.7 × 10^−5^, slightly smaller but similar in magnitude to the angle distribution predicted by the simulation.

## 3 Discussion

As long-range cellular projections like airinemes continue to be discovered in multicellular systems, their mathematical characterization will become increasingly valuable, mirroring the mathematical characterization of diffusion-mediated cell-cell signals. We have measured the *in situ* shape of airinemes, and find agreeable fit to a finite, unobstructed persistent random walk model, rather than Levy or diffusion-like motion. The mean-square curvature, or equivalently the directional persistent length, is close to that which allows optimal search efficiency for target cells. Since airineme tip motion is driven by macrophages, our results have implications for macrophage cell motility, which is relevant in other macrophage-dependent processes like wound healing and infection [40, 41].

Besides search success probability and directional information, there are other objective functions that airinemes and other cell protrusions could be optimized for. One obvious candidate is the efficient transport of signaling molecules after contact has been established [42, 43], which might decrease monotonically with protrusion length, and would therefore favor straight protrusions. Notably, nanotubes in cancer cells [26] are strikingly straight compared to airinemes, which might suggest a different balance between objectives. In the future it would be intriguing to compare all known non-canonical protrusions in light of the three performance objectives, and others.

The cell-cell interactions mediated by airinemes contribute to large-scale pattern formation in zebrafish, a subject of previous mathematical modeling [27, 44, 45]. Our results provide a contact probability per airineme, setting an upper-bound on the ability of cells to communicate via this modality, which is itself a function of cell density (related to *d*_targ_ in our notation). Thus our results may inform future pattern formation models. In the reciprocal direction, these models may provide information about the distribution of target cells, which may significantly affect search efficiencies.

## 4 Materials and Methods

### 4.1 Zebrafish husbandry and maintenance

Adult zebrafish were maintained at 28.5°C on a 16h:8h light:dark cycle. Fish stocks of *Tg*(*tyrp1b:palmmCherry*)*^wp.rt11^* [47] were used. Embryos were collected in E3 medium (5.0mM NaCl, 0.17mM KCl, 0.33mM CaCl2, 0.33mM MgCl2·6H2O, adjusted to pH7.2-7.4) in Petri dishes by in vitro fertilization as described in Westerfield with modifications [48]. Unfertilized and dead embryos were removed 5 hours (hpf) and 1-day post-fertilization (dpf). Fertilized embryos were kept in E3 medium at 28.5°C until 5dpf, at which time they were introduced to the main system until they were ready for downstream procedures. All animal work in this study was conducted with the approval of the University of California Irvine Institutional Animal Care and Use Committee (Protocol #AUP-19-043) in accordance with institutional and federal guidelines for the ethical use of animals.

### 4.2 Time-Lapse and static Imaging

The transgenic embryos, *Tg*(*tyrplb:palmmCherry*), were injected with the construct drive membrane bound EGFP under the aox5 promoter to visualize airinemes in xanthophore lineages and melanophores [30]. Zebrafish larvae of 7.5 SSL were staged following [49] prior to explant preparation for ex vivo imaging of pigment cells in their native tissue environment as described by [50, 51]. Time-lapse images, acquired at 5-min intervals for 12hr, and static images were taken at 40x (water-emulsion objective) on a Leica SP8 confocal microscope with resonant scanner.

### 4.3 Model definitions, simulation and analysis

In the finite-length persistent random walk model (PRW), the position of the airineme tip at time *t* is given by

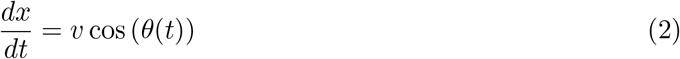

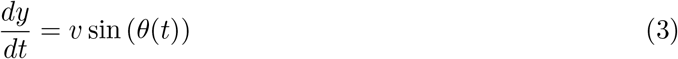

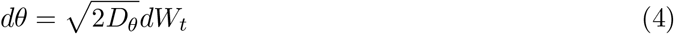

where *W_t_* is a Wiener process, and *D_θ_* is related to the directional persistence length *l_p_* in 2d by *D_θ_* = *v*/2*l_p_*.

The MSD for PRWs is [32, 33]

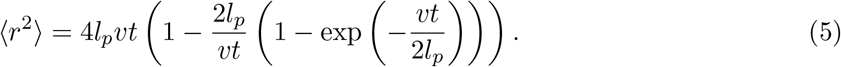

Our persistent random walk model is finite-length, meaning we assume the airinemes extend only up to *l*_max_ = 250*μm*. This assumption yields a final length distribution (Fig. S4) consistent with the observed distribution [30].

To simulate this model, we use an Euler-Maruyama scheme with timestep Δ*t* ≪ 1/*D_θ_*, implemented in Matlab (The MathWorks). To validate these simulations, at two limits of *D_θ_*, search contact probabilities can be solved analytically (Fig. 3AB filled circles). First, the straight limit *D_θ_* → 0. Suppose an airineme searches for the target cell centered at (0,0) with radius r_targ_, and the airineme emanates from a source at (*r*_targ_ + *d*_targ_, 0). Let *ϕ* be the angle between the hitting point on the target cell and the center line. Then,

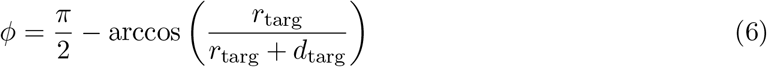

and *P*_contact_ = *ϕ/π*. At the other limit, *l_P_* ≪ *d*_targ_, the PRW is approximately equivalent to diffusion with coefficient *D* = *v*^2^/2*D_θ_*. For a finite time 0 < *t* < *l*_max_/*v* diffusive search process, the probability of hitting the target cell is *P*_contact_ = 1 – *S*(*r, t*) where *S*(*r, t*) denotes the survival probability, which evolves according to

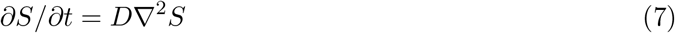

with *S*(*r, t*) = 0 on the surface of the target cell. We solve this PDE and display results in Fig. 3AB blue circles. For these validations, the *D_θ_* values were chosen to fit the blue circles onto the plot.

### 4.4 Image analysis and model fitting

In order to estimate uncertainty in our analysis method, we used model convolution (Fig. S3, [35]). Specifically, we first measured the experimental signal-to-noise ratio and point-spread function. We then simulated airinemes with a ground-truth curvature value, and convoluted the simulated images with a Gaussian kernel with the signal-to-noise ratio and point-spread function measured from experimental data. Since there is a manual step in this analysis pipeline, independent analyses by five people were performed on both simulated data and experimental data. For simulated data, the difference between simulated and estimated *D_θ_* was less than 7% in all cases and usually ~ 2%.

### 4.5 Directional information

To measure the directional information that, stochastically, an airineme provides its target cell, we use the Fisher information,

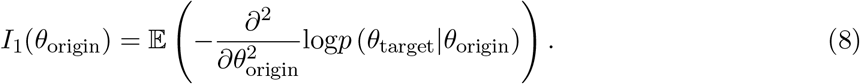

This quantity can be intuitively understood by noting that, for Gaussian distributions, Fisher Information is the inverse of the variance. So, high variance implies low information and low variance implies high information. If multiple independent and identically distributed airinemes provide information to the target cell, then the probability densities of each airineme multiply, and the Fisher Information is *I*(*θ*_origin_) = *n_hit_* · *I*_1_(*θ*_origin_) where is the number of successful attempts, which is proportional to *P*_contact_. Therefore, we define the modified Fisher Information

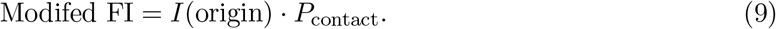

With this modification, a target cell that receives almost no airinemes will score low in directional information.

### 4.6 Experimental measurement of directional information

Images capturing incidences of airinemes with membrane-bound vesicles extended from xanthoblasts, stabilizing on melanophores were captured and imported into ImageJ for angle analysis between a. Originating point of airinemes on xanthoblasts, b. Center of target cells (i.e., melanophores), and c. Docking site of airineme vesicles on target cells. Two intersecting lines were drawn as follows: 1. Connect the originating point of an airineme on a xanthoblast with the center of the target melanophore to draw the first line, and 2. Connect the center of the target melanophore to the docking site of the airineme vesicle on the target melanophore to draw the second line. The angle between the three points connected by the two intersecting lines was then generated automatically with the angle tool in ImageJ. Coordinates of each point and the corresponding angle were recorded by ImageJ and exported to an Excel worksheet for further analysis. Each angle was assigned a ± sign in the 180-degree system based on the relative location of the three points at the time of the airineme incident. The 0-degree line was defined as the line passing through the center of the target melanophore. Thus, a positive angle was assigned when the originating point of an airineme on a xanthoblast lies on the 0-degree line with the docking site of airineme vesicles on the target melanophore lies above the 0-degree line, and vice versa.

## 5 Acknowlegdements

We thank Sean Lawley (U Utah), Jay Newby (U Alberta), and Yoichiro Mori (U Penn) for valuable discussion. We acknowledge support from NSF CAREER award DMS-1454739 to JA, NIH R35GM142791 to DSE, NSF grant DMS 1763272 and two grants from the Simons Foundation (594598, QN and Math+X grant to the University of Pennsylvania).

## 6 Author contributions

SP designed and performed the image analysis, developed model, simulation and analysis, and prepared the manuscript. HK designed the project and models, and developed model, simulation and analysis. YW performed image analysis. JA and DSE designed the project and prepared the manuscript.

## 7 Supplemental Figures

**Figure S1:**
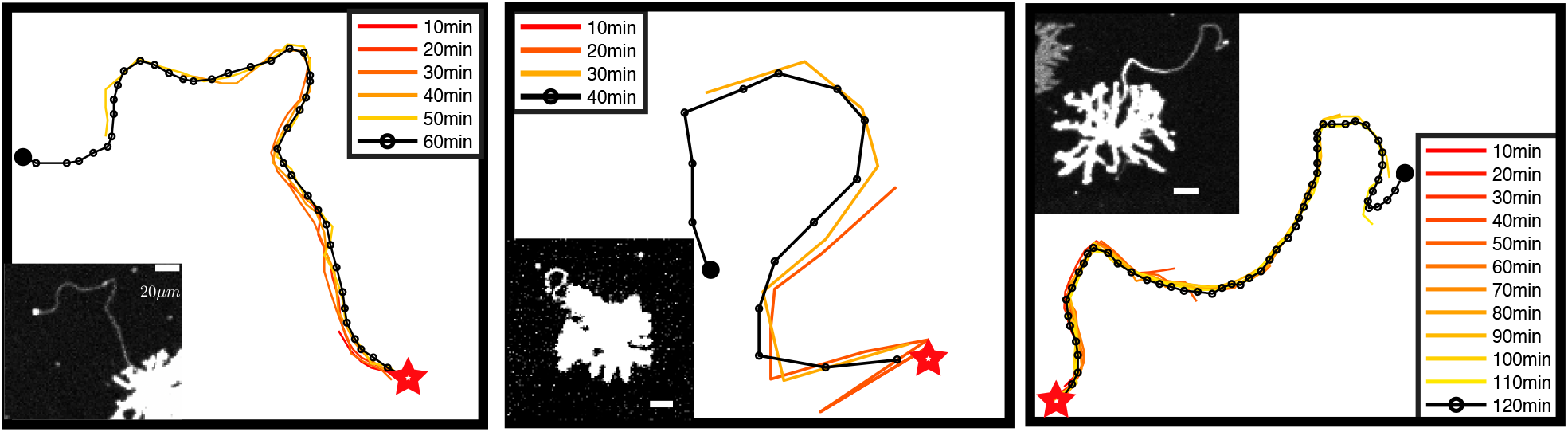
Time evolution and final state are self-consistent. Each panel shows the examination of a single representative airineme as it emerges from the source cell at *t* = 0, large star. Every 10 minutes, the shape is tracked at discrete points as it grows away from the source cell. The micrograph of the final length is shown as an inset (only the channel labeling the source cell is shown, so the target cells are not visible). Scale bar is in 20*μm*. We computed root mean squared curvature of both the time series tip location and of just the final state for the airinemes shown here. These two methods yield a difference, on average, of 0.7%. Thus, we conclude airineme shape does not change significantly throughout its extension.

**Figure S2:**
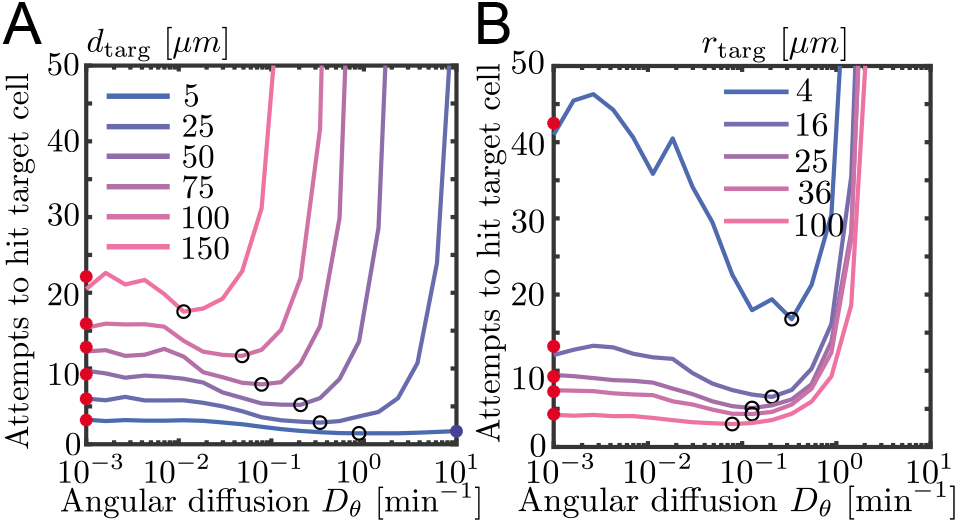
Simulated contact success shown as number of attempts instead of probability of contact. Same as Fig. 3AB in main text, but showing contact success as mean number of attempts (= l/*p*_contact_).

**Figure S3:**
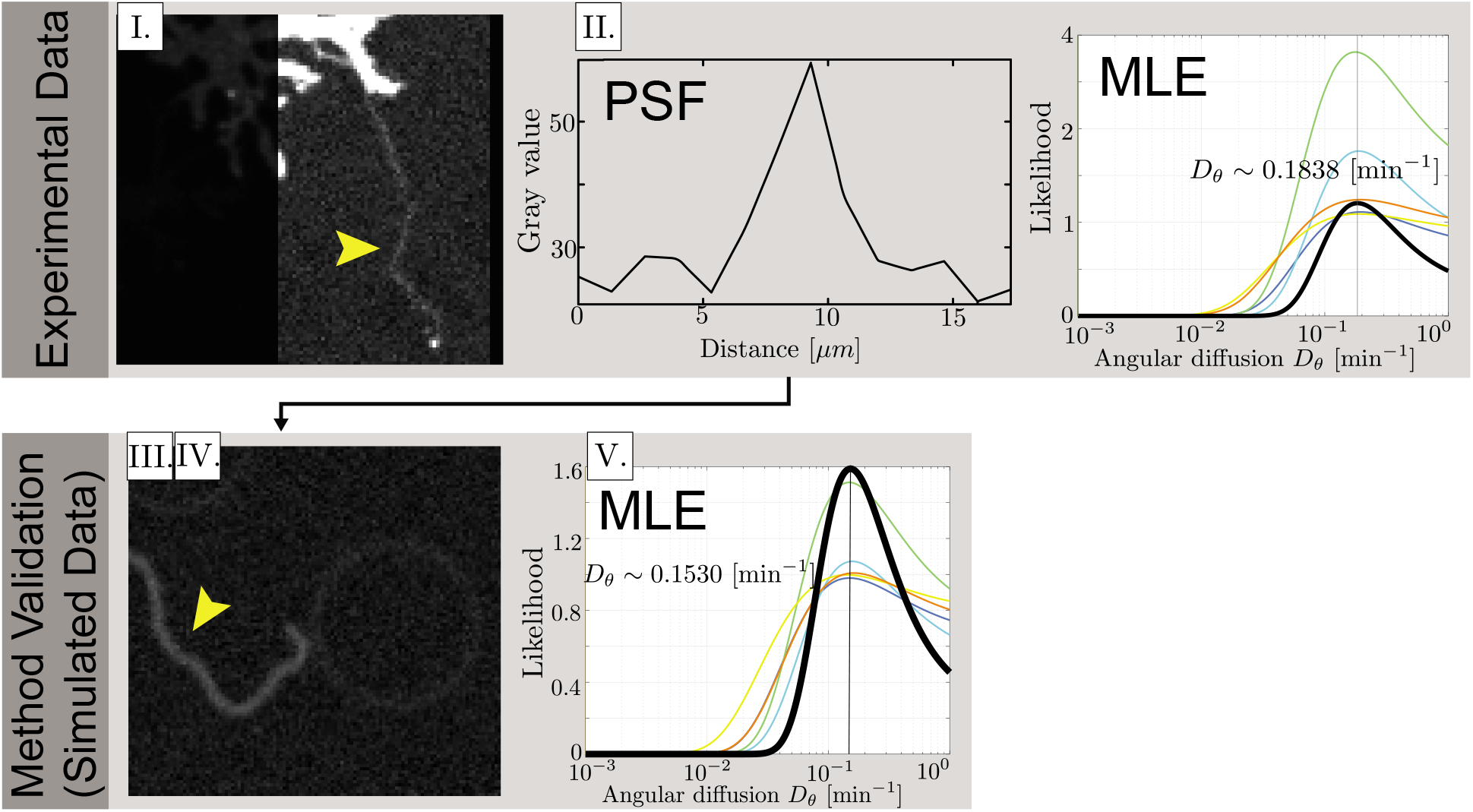
Experimental data analysis method validation and data collection. I. Fully extended airineme micrograph image. Arrow indicates the airineme. II. Extraction of the average noise level from image. III. Images of simulated airinemes are convoluted to have the same statistical noise properties of the experimental image. We use a “ground truth” *D_θ_* ~ 0.1525min^−1^. IV. Five people used a manual image analysis pipeline to locate points along airinemes. From these points, curvatures and autocorrelation functions were computed, and *D_θ_* likelihoods extracted. Curvature of simulated airineme images and likelihood distributions are shown as colored curves. Black curve shows the likelihood distribution for combined data. V. The maximum likelihood *D_θ_* value was close to the simulation input *D_θ_*, with typically 2% error. The most erroneous (yellow curve) was 7% and corresponds to the manual entry of the corresponding author, J.A. (Top right) Likelihoods of *D_θ_* estimates from experimental images from 5 people (colored curves) and combined (black curve).

**Figure S4:**
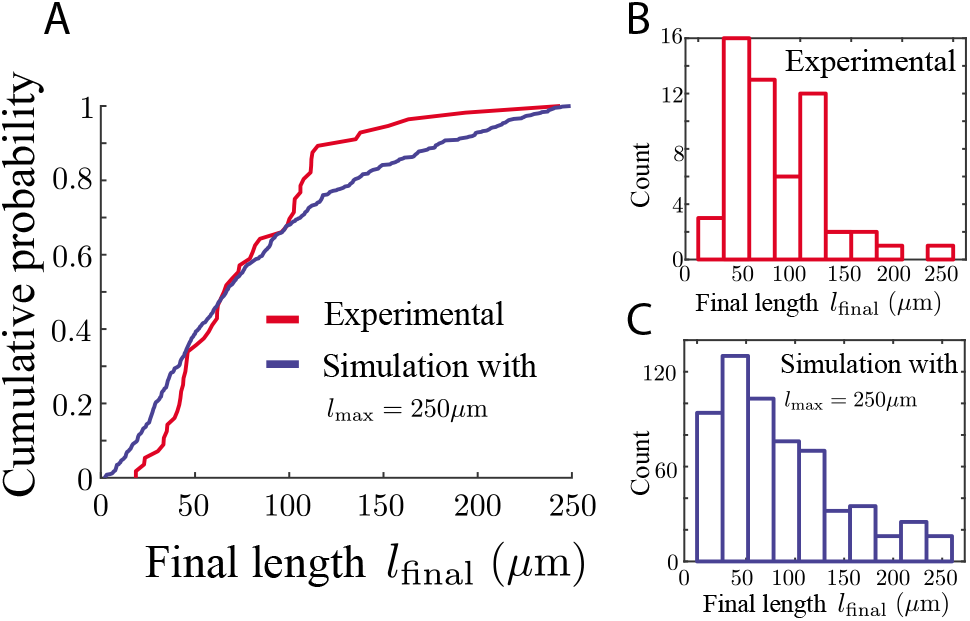
Agreement with final lengths of airinemes. For the model parameter *l*_max_, which is the length at which an airineme stops searching if it has not yet found a target, we wondered whether a single number would be appropriate or whether we needed to use a distribution to capture the variability of airineme lengths. We measure the experimental quantity *l*_final_, which is the length of airinemes when they stop growing, whether or not they have made contact with a target cell, shown in red in A,B (*N* = 56 airinemes). We run simulations in which target cells are randomly placed according to 2d Poisson statistics with mean distance *d*_targ_ = 50*μm* and measured their length distribution when they stop growing *l*_final_, shown in blue in A,C (again, whether or not they have made contact with a target). We find that the distributions for experimental data and simulated data agree qualitatively when *l*_max_ = 250*μm*. Note the variability in both experimental and simulated distributions is of similar magnitude, suggesting the randomness in observed airineme length arises from the randomness in target cell placement.

**Figure S5:**
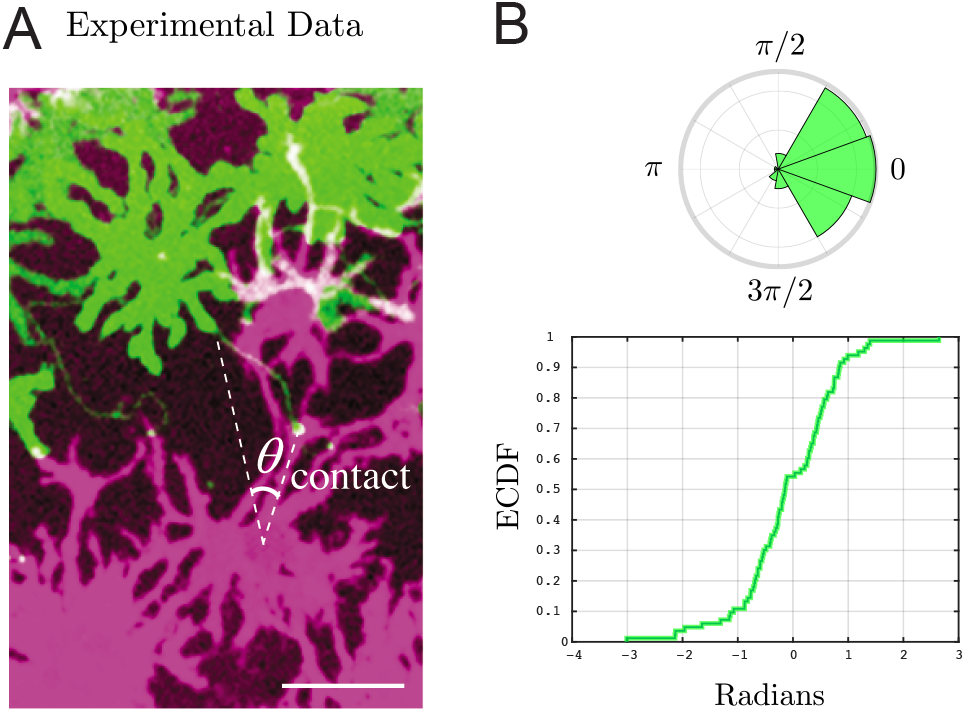
Experimental measurement of contact angle and directional information. (A) Representative image of source cell (green), airineme (light green, touching two white lines), and target cell (purple). To estimate the contact angle distribution, we draw a line from the source of the airineme to the center of the nucleus of the target cell, then another line from here to the airineme contact point. Scale bar is 50*μm*. (B) The angle distribution of contact angles shown in an angular histogram (top) and empirical cumulative distribution (bottom). From these, compute (unmodified) Fisher Information of of FI 3.93 × 10^−4^rad^−2^, and modified Fisher Information to be 5.7 × 10^−5^rad^−2^ from Eq. 9. Data from *N* = 83 airinemes.

